# Genetic variation in host selectivity and adaptive strain enrichment within legume-rhizobia symbiosis: processes are host-dependent, far from perfect, and correlate with nodule morphology

**DOI:** 10.1101/2025.06.16.659998

**Authors:** Liana Burghardt, Patrick Sydow, Jeremy Sutherland, Brendan Epstein, Peter Tiffin

## Abstract

Mutualism breakdown can be prevented if partner species preferentially select and reward partners that provide greater benefit. We examined these two components using the legume *Medicago truncatula* and its nitrogen-fixing symbiont *Sinorhizobium meliloti*. First, we reanalyzed data from 202 accessions to show significant genetic variation in the capacity of *Medicago* to restrict strain diversity, finding that hosts with shorter nodules were more selective. A genome-wide association study on host selectivity identified genes including the hormone leginsulin, pectin degradation, multidrug and toxic compound efflux, Zinc transport, and DNA methylation. Second, we used two well-studied *Medicago* genotypes with contrasting nodule morphologies to assess the effectiveness of adaptive enrichment mechanisms by sampling the relative frequencies of rhizobial strains in pools of small nodules (indicating a lack of host investment) compared to large nodules (indicating increased host investment) and pairing these results with previous single-strain assessments of strain benefits to hosts. While both hosts enriched beneficial strains in large nodules, the host that formed larger and more variably sized nodules and thus had greater ’potential’ to increase rhizobial populations was less effective. Our findings reveal that host genetic variation affects strain selectivity and suggest that nodule traits warrant attention when exploring mutualism evolution.

## INTRODUCTION

Mutualisms occur when two species derive fitness benefits from interacting with each other (Bronstein, 2015). Mutualism’s stability depends on mechanisms limiting the spread of genotypes that do not benefit their partners (Sachs et al., 2004). These partner choice mechanisms may operate before resources are exchanged via, for instance, signaling and recognition (Bull and Rice, 1991), or after partners start to exchange benefits (e.g., (Denison, 2000; Kiers et al., 2003; Bever et al., 2009)). Post-exchange mechanisms, such as sanctions and rewards, both involve the host altering the fitness of symbionts but differ in that sanctions block the reproduction of less beneficial symbionts, whereas rewards direct resources to beneficial symbionts (Bever et al., 2009; Bever, 2015). Despite the centrality of partner choice mechanisms to mutualism evolution, little is known about natural genetic variation in these traits within host species (Porter et al., 2024). When and which of these mechanisms hosts use is important because they have different effects on maintaining variation in the symbiont quality (Bever, 2015). Here, we characterize natural genetic variation in host selectivity and adaptive enrichment in the model mutualism between *Medicago truncatula* and *Sinorhizobium meliloti*.

Legumes and rhizobia form a resource-based mutualism that is a model for investigating the genetics and evolution of plant-bacteria symbiosis (Sachs et al., 2004, 2018; Oldroyd et al., 2011; Friesen, 2012; Burghardt et al., 2018). This mutualism is formed when soil-inhabiting rhizobia invade a legume root, triggering the plant to develop a nodule. Inside the nodule, rhizobia convert atmospheric N_2_ to a plant-usable form. In exchange, the plant provides the bacteria with carbon compounds, which, along with the environment inside the nodule, support a high rate of bacterial reproduction (Sprent et al., 1987); hundreds of thousands of bacteria can emerge from a nodule formed by a single bacterium (e.g. (Sachs et al., 2010)). Although a single bacterium invades most nodules, individual plants, which can form hundreds of nodules, can form nodules with many different rhizobial strains. It is vital to study mutualism maintenance in horizontally transmitted mutualisms like this because they are considered particularly susceptible to mutualism breakdown as rhizobia spend many generations living in soil and rhizosphere, between inhabiting nodules (Wilkinson and Sherratt, 2001).

Pre-nodulation selection mechanisms may be less effective at filtering for beneficial partners than post-nodulation mechanisms because hosts form associations before receiving feedback on whether the rhizobia they associate with are beneficial (Wang et al., 2012). However, once nodules have formed, host species may allocate resources based on the resources being provided by individual nodules, thereby differentially allocating resources to protect themselves from investing in less-beneficial rhizobia (Porter et al., 2024). For example, by reducing N_2_ availability to nodules, researchers have shown that hosts can withhold resources from nodules (i.e., sanctions) when no N-fixation occurs (Denison, 2000; Kiers et al., 2003). However, it is less clear if sanctions are effective when N-fixation is reduced rather than eliminated (Kiers et al., 2006). Nevertheless, several studies suggest enrichment can occur after nodules form if there is a substantial difference in N-fixing capacity between strains (Quides et al., 2017, 2021; Regus et al., 2017; Burghardt et al., 2018; Wendlandt et al., 2019; Westhoek et al., 2021; Montoya et al., 2023; Underwood et al., 2024). One theme emerging from these studies is that hosts may be able to enrich strains by funneling resources to increase the size of nodules with helpful strains and limiting resources to nodules inhabited by less helpful strains (Porter et al., 2024). These findings suggest that nodule size traits and morphological variation that have received little empirical attention may provide new avenues for studying partner choice mechanisms.

Nodule development differs among legume species, with some forming nodules with determinate meristems and others forming nodules with indeterminate meristems (Brewin, 1991; Łotocka et al., 2012). This difference can have consequences for rhizobial populations. In species with determinate nodule development, nodules have a transient meristem that is active during early stages of development, creating spherical nodules with little morphological variation beyond size. In contrast, in species that form indeterminate nodules, the nodule meristem keeps dividing, causing ongoing nodule growth and elongation, increasing the zone of N-fixation, and increasing the population size of the rhizobia in the nodules. The meristem of indeterminate nodules can also branch, increasing the habitat space for rhizobial populations (Brewin, 1991). Here, we examine the relationship between nodule morphology and host selectivity. Our initial prediction was that host genotypes with larger nodules and higher variance in nodule size have a greater potential to enrich helpful strains by boosting populations in large nodules.

We took a two-pronged approach to evaluate i) genetic variation among Medicago hosts in their ability to select or adaptively reward more beneficial rhizobia strains and ii) the extent to which nodule morphology is associated with differential rewards (Figure 1). First, to examine if there is natural genetic variation for host selectivity, we extended the analysis of a previously published dataset including 202 *M. truncatula* accessions (Epstein et al. 2023) by accounting for differences in nodule number across samples and expanding the nodule traits examined. Second, we collected data on two well-studied *Medicago* accessions with intermediate host selectivity and contrasting nodule morphology patterns to assess the efficacy of pre- and post-nodulation selection mechanisms. We grew each host with a mixture of 68 naturally occurring rhizobial isolates. To assess how rewards vary across contrasting nodulation patterns (high vs. low size variation), we sampled rhizobial strains in pools of large nodules (representing increased host investment) vs. small nodules (lack of investment after nodule formation). We show that host selection of beneficial rhizobia occurred both before and after nodules formed, and host selectivity is associated with shorter and more numerous nodules, and enrichment was more effective in the host with shorter, more numerous nodules.

**Figure 1:**
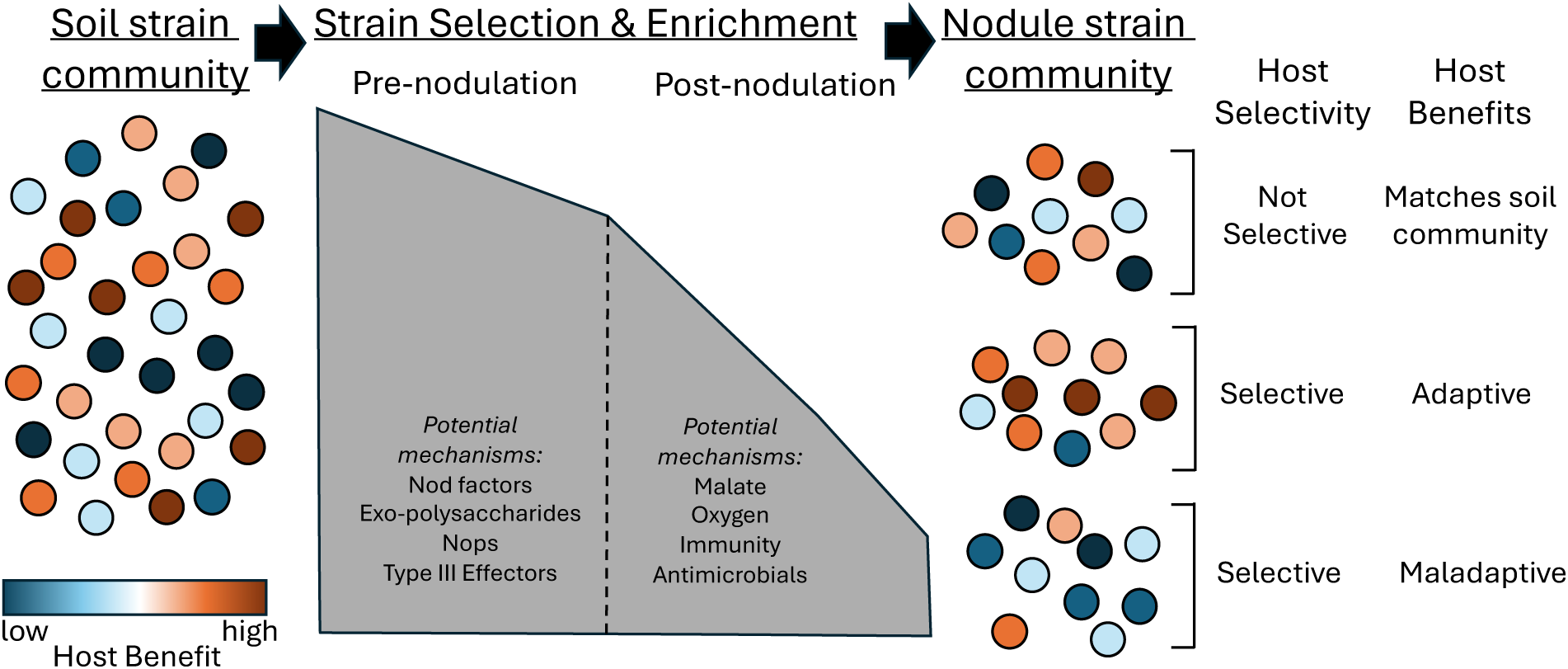
Host selectivity and adaptive strain enrichment in legume nodules. Determining if hosts can choose partners, when (before or after nodules form), and via what mechanisms (e.g., recognition, resource allocation) remains an essential question in mutualism research (Porter et. al, 2024). Here, we examine host genetic variation in two complementary ways. First, we examine host selectivity for rhizobia in 202 accessions of *Medicago truncatula.* By selectivity, we refer to hosts associating with a non-random sample of rhizoiba in the innoculm. Regardless of how beneficial those strains are. Second, we assess two specific hosts’ ability to adaptively enrich the fitness of beneficial strains by increasing the size of nodules and, thus, the population sizes of those strains.

## MATERIALS AND METHODS

### Genetic variation in host selectivity and nodule morphology

To determine the breadth of genetic variation in nodule morphology and host selectivity, we expanded the analysis of a study that grew 202 accessions of *Medicago truncatula* (the HapMap panel) with a mixture of 89 strains of *Sinorhizobium meliloti* (Epstein et al., 2023). Epstein et al. (2023) estimated strain diversity among the nodules on each plant based on the frequency of each strain in the nodules. However, nodule number was positively correlated with strain diversity (R=0.26, *p* < 0.001). The among-accession differences in nodule number may, therefore, obscure mechanisms that more directly affect host selectivity of differential rewards. To account for the effect of nodule number, we calculated a metric of host selectivity that controlled for nodule sampling via simulations of the nodule community. In our simulations, the number of nodules sampled from each plant determined the number of random draws from the initial 89-strain community. In this model of assembly, we assumed nodules contained a single strain of rhizobia as coinfection is rare (3-20% of nodules)(Daubech et al., 2017)(Figure S1). Simulations were completed in R, and the probability of drawing a strain was equal to the relative frequency of the strain in the initial inoculum. We calculated host selectivity as H_s_ = H_e_ - H_o_ where H_s_ = host selectivity, H_e_ = expected Shannon diversity (mean H from the 10 simulations) and H_o_ = observed Shannon diversity. Shannon diversity (H) was calculated with the vegan package version 2.6-4 (Oksanen, 2022). Host selectivity values above zero indicate a host with a less diverse strain community than expected by chance, while values below zero indicate a more diverse community. To determine the major ways nodule morphology varies across host genotypes, we first summarized traits measured on individual nodules (e.g., area) by pot. We then completed a redundancy analysis constraining 16 traits describing nodule morphology (Table S1) by host genotype and experimental block. Pearson correlations between genotypic means of traits were also calculated and visualized using the R package ‘corrplot’. Clusters of nodules traits were then identified using Ward clustering criterion on squared dissimilarities of genotypic means using the hclust() function in R.

### Associating Medicago genetic variation with host selectivity

To determine host genomic regions associated with the host selectivity, we conducted a Genome-wide Association Study (GWAS) using a linear mixed model (LMM) implemented with the -lmm option in GEMMA (v0.98.3) (Zhou and Stephens, 2012). We obtained the SNP data for *Medicago truncatula* A17 from the Legume Information System (legumeinfo.org, (Dash et al., 2016). The SNP data were initially filtered using bcftools v1.17 (Li, 2011) to include only biallelic SNPs with a minor allele frequency greater than 0.05. We then filtered the remaining SNPs in plink v1.90b6.21 (Purcell et al., 2007) to a missingness threshold of 0.2. We included a standardized K-matrix (*r^2^*>= 0.3) for the GWAS to reduce the effects of unequal relatedness among accessions. *P*-values for each SNP were calculated with a likelihood ratio test, using a *p*-value cut-off of 1×10^-7.^ Lastly, we used bedtools to find annotated genes closest to each candidate SNP in the Mt5.0 reference genome (Table S2).

### Associating host selectivity with nodule morphology characteristics

To identify relationships between nodule morphology and host selectivity, we first calculated Pearson correlation coefficients between genotypic means of host selectivity and all nodule traits (Table S1). Measures of variance for nodule traits within individuals were excluded due to high correlations with values of nodule trait means, even after scaling by trait means (R^2^ > 0.6). To identify which group or cluster of traits was most associated with host selectivity, we used Elastic Net regression in R (glmnet package version 4.1-7) (Friedman et al., 2010) with 15 traits describing nodule morphologies as predictors (Table S1). Elastic net regression is a machine learning shrinkage algorithm that balances the strict (L1) and relaxed (L2) penalties of LASSO and ridge regression to identify groups of correlated parameters (i.e., traits) that form a linear regression model with the smallest error. The number of parameters is determined by the size of their coefficients, with the tuning parameter (λ) determining the magnitude of the shrinkage penalty. Here, λ was determined via ten-fold cross-validation after the traits were centered and scaled, with λ assessed using the mean squared error (Figure S4). To identify which single trait is most associated with host selectivity, univariate linear models using the genotypic means of traits selected by elastic net regression were compared based on their significance and percent variance explained (R^²^) (Table S3). Once identified, we further characterized the nodule trait-host selectivity relationship by creating a quadratic polynomial regression model with reduced error.

### Select and Resequence experiment to examine strain enrichment in small vs. large nodules

We chose two divergent host genotypes that represented the extremes of the variation in nodule morphology, and have intermediate host selectivity values, A17 and R108, which are also genetic models in this system.

To estimate the fitness enrichment that individual rhizobia strains receive from these hosts, we inoculated both accessions with a 68-strain community of *S meliloti*. Most strains were from 101 strains obtained from the US Department of Agriculture (USDA) strain collection and used to investigate host x strain interactions in nodulation (Burghardt et al., 2018, 2020). Notably, this strain community represents an independent set of strains from those used in the previous experiment.

We grew six replicates of each of the two host accessions. Each replicate comprised 10-12 plants grown in a 1L pot filled with Sunshine Mix #5. Before planting, we disrupted seed coats with a razor blade, incubated the seeds on wet filter paper for two days at 4°C, and then left the seeds at room temperature overnight to germinate before planting. The rhizobia inoculum was formed by mixing equal volumes of 68 single-strain cultures grown in TY media for 72 hours at 30°C with shaking at 200 rpm.

When seedlings were three days old, we inoculated each pot with approximately 10^8^ rhizobia from the mixed inoculum diluted in 10 ml of 0.85% NaCl solution. Thereafter, we watered pots as needed and fertilized them once a week with 150 mL of N-free fertilizer (Bucciarelli et al., 2006). Ten weeks after planting, we harvested the plants. At harvest, we removed nodules from each pot and divided them into three size classes (small, medium, and large) based on visual size estimates; for A17, small and large pools from each replicate contained the same number of nodules. However, for R108 plants, the small nodule pools contained 1.5-2x as many nodules for three of the six pots as the paired large nodule pools. This was because R108 formed far fewer nodules, with many massive ones, and we encountered limitations in our processing volumes; however, this did not reduce strain diversity (Figures S5-6).

We characterized the small and large nodule pools in three ways: 1) To estimate the number of rhizobia released from a single representative nodule in each size class, we counted the colony-forming units (CFUs) from serial dilutions of a randomly chosen nodule from each size class from each replicate pot. We sterilized the surface of each nodule in 20% bleach for 90 seconds, added 1 ml of sterile 0.85% by volume NaCl solution, and crushed individual nodules with a pestle. We then used the nodule slurry to create three dilution series and plated 100 μL of each series on two replicate TY plates. After 48-72 hours at 30°C, we counted the colonies on each plate. 2) To assess nodule characteristics in each group, we counted lobes, measured nodule length (mm) of 10 representative nodules from each size class, and counted and weighed each pool of nodules. 3) To determine the frequency of each rhizobial isolate in each nodule pool, we followed Burghardt et al.’s Select and Resequence method (2018). Briefly, we surface-sterilized each nodule pool in 20% bleach for 90 seconds and crushed the nodules with a pestle in a 2 mL tube containing 1 mL of sterile 0.85% NaCl solution. We centrifuged the slurry at 200xg for 10 min to pellet heavier plant tissue and bacteriods and create a supernatant enriched for undifferentiated rhizobial bacteria. We then pelleted undifferentiated rhizobia via fast centrifugation (20,000xg for 4 min), removed the supernatant, and froze the pellets at −20°C. Following the manufacturer’s instructions, we extracted DNA using Qiagen Plant DNEasy kits.

### Sequencing and analysis of large and small nodule pools

The University of Minnesota Genomics Core sequenced DNA from all samples with an Illumina HiSeq 2500, generating paired-end 125 bp reads. We removed adapters and low-quality sequences by running TrimGalore! (v0.4.1; github.com/FelixKrueger/TrimGalore) with the following settings: –quality 30, –stringency 3, –length 100, –length_1 101, –length_2 101 −e 0.1. We mapped the trimmed reads to the *S. meliloti* USDA1106 reference genome using bwa mem v0.7.17 (Li and Durbin, 2010) with default options, and then employed HARP (Kessner et al., 2013) to estimate strain frequencies. SNPs segregating among the 68 strains were previously identified by sequencing each strain using Illumina (Burghardt et al., 2022). The Illumina reads were trimmed with TrimGalore! (v0.4.1, settings −e 0.1, –quality 30, –stringency 3, – length 126 or 70, –length_1 and –length_2 127 or 71; shorter lengths were used for Illumina GAIIx data, longer lengths were used for Illumina HiSeq 2500 data), aligned to USDA1106 with bwa mem (v0.7.17), and the SNPs identified with freebayes (v1.2.0-2), and filtering out sites with QUAL scores < 20.

### Statistical analysis of nodule morphologies, strain communities, strain enrichment, and host benefits

To assess nodule morphological (length, branching, weight) and population size differences across hosts and nodule size pools, we ran linear models (ANOVA) in R on log-transformed values for nodule length, lobes, and population sizes: lm (Trait ∼ host*pool size). We measured strain frequencies in the initial community (median=0.0092, 5%-95% quantile: 0.0054-0.0154).) and large and small nodule pools for each replicate pot. We calculated strain relative fitness = log_2_(selected frequency/initial frequency) to control for slight differences in strain frequencies in the inoculum and normalize the data. To quantify and visualize multivariate shifts in strain fitness and nodule morphology across nodule pools, we conducted a Redundancy Analysis using the ‘rda’ function in the ‘vegan’ R package (Oksanen et al., 2017), which fits a multivariate linear regression to centered and scaled data and uses PCA to decompose the principal axes of variation in the fitted parameters. The adjusted R^2^ of the RDA model provides an estimate of the proportion of variance in relative fitness explained by the model predictor(s) and, when paired with a PERMANOVA (‘anova’ function, 999 permutations), provides an estimate of the proportion of variance in relative fitness explained by each predictor. To assess shifts in host benefits of strains in large and small pools, we calculated the proportion of the potential benefit each host captured in each replicate of small vs large nodule pools, inferred from the benefit each strain provides to the host. The benefits each strain provides to hosts were derived from published data on the size of plants when inoculated with single strains under nitrogen-free conditions (Burghardt et al., 2018). Using ANOVA, we tested for 1) enrichment above random assembly based on the initial inoculum and 2) differential enrichment across nodule sizes within a host. Lastly, we sought to determine the relationship between host benefit and strain rewards by using strain representation in small vs large nodule pools as a signal of host nodule investment. To measure differences in *relative strain enrichment in large nodules*, we summed the strain frequency in small and large nodules for each replicate pot.

Then we divided the relative frequency in large nodules by the total value and took the median value across pots. A value of 0.50 means that the strain is equally represented in small and large nodules. 0.99 means 99% of the total strain frequency was found in large nodules, and 0.01 means only 1% was in large nodules. To determine whether the strains enriched in large nodules are more beneficial to hosts, we ran a linear model to see if *host benefits provided by each strain* predicted strain enrichment in large nodules (strain enrichment ∼ host biomass).

## RESULTS

### Host selectivity was strongly heritable and associated with maintaining shorter nodules

After using nodule assembly simulations to remove the confounding effect of nodule number on strain diversity, our estimates of host selectivity across the Medicago GWAS panel of 202 accessions ranged from −0.22 to 0.95, with a median of 0.14 (Figure 2a). Positive host selectivity values mean that host nodule pools were less diverse than expected if the host randomly formed associations with the 89 potential rhizobial strains. The effect of host genotype on selectivity was strong, explaining 42% of the variation (*p*<0.001). Most (94%) hosts were more selective than expected by chance. While most were weakly selective there was a long tail of more highly selective accessions. These results suggest that most hosts weakly filter the strain diversity found in the initial community, which is composed of all 68 strains; however, segregating natural genetic variation exists, making hosts more selective.

**Figure 2:**
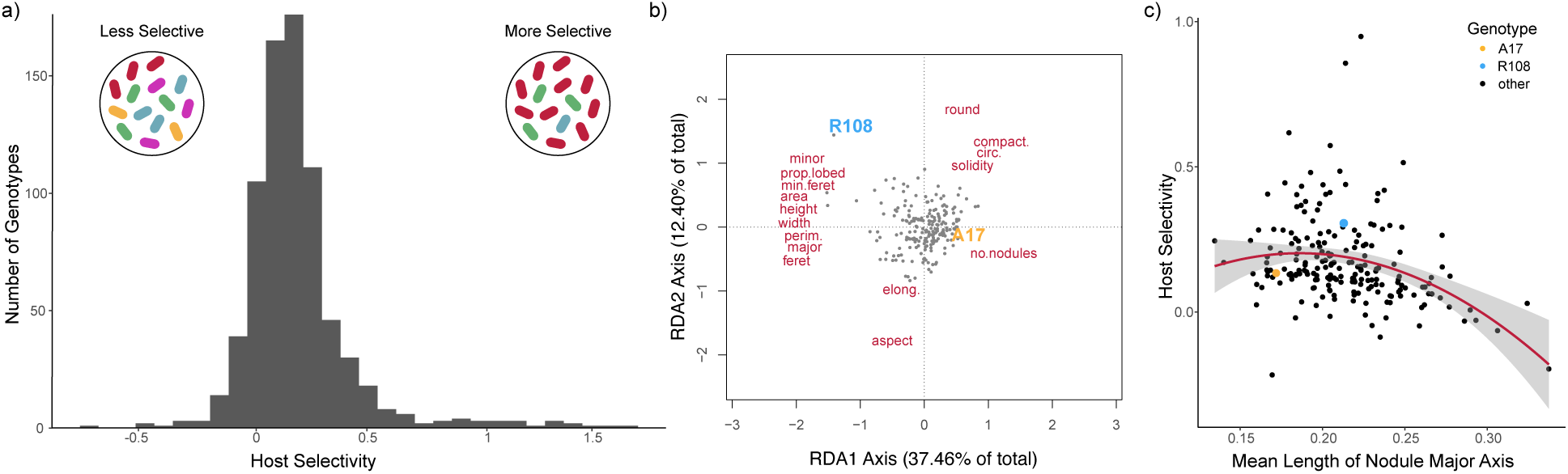
Host selectivity and nodule morphology. a) For 202 accessions of *Medicago truncatula,* we calculated host selectivity while controlling for nodule number differences among plants, and b) quantified nodule morphology variation in nodule pools (16 trait means, red text, are visualized here with a redundancy analysis, RDA). Dots represent host accessions. We used a combination of elastic net regression followed by univariate regression modeling to determine the mean length of the major axis of nodules to be the best predictor (Figure S4; Table S3) of host selectivity (c). Hosts used for our follow-up experiment are highlighted in blue (R108) and gold (A17). Lastly, we used GWAS to identify host candidate genes associated with host selectivity (Table S2).

We performed a GWAS to identify host gene regions associated with host selectivity, which had a strong genetic basis (broad sense H^2^ = 0.465). The top ten associations had *P* values less than 10^-7^, with effect sizes ranging from 0.09 to 0.16. These SNPs tag several genes, many with known effects on symbiotic interactions, including leginsulin 37, MtZIP11 (a Zinc-Iron permease), a pectate lyase gene, MtMATE38 (a multidrug & toxic compound extrusion protein), and Spt4, which encodes a putative transcription initiation factor (Figure S2, Table S2).

Lastly, we explored if any nodule morphology traits were associated with host selectivity (Figure 2b, Table S1). Nodule morphological traits fell into five clusters of highly correlated traits (all r > 0.81, Figure S3). Overall, hosts that formed smaller, shorter, less elongated, more compact, and more numerous nodules tended to be more selective. Using a combination of elastic net regression and univariate regression modeling, we found that smaller and shorter nodules were associated with greater host selectivity, with the mean major axis length, width, and area of nodules each explaining a modest 8-9% of variation in host selectivity (Figure 2c; Table S3; Figure S4). Post hoc polynomial regressions of significant univariate predictors indicated an intermediate optimum for major axis length, which explained ∼12% of variation in host selectivity, even when outliers are removed, suggesting that hosts that form very long or short nodules tend to be less selective. This finding suggests that host control of nodule shape may be more important than overall differences in nodule size for host selectivity.

However, our results also suggest a limited relationship between nodule morphology traits and host selectivity.

### Host genotypes differed in their ability to enrich populations of beneficial strains

With the broad approach above, we cannot determine whether selective hosts are better or worse at increasing the fitness of beneficial strains of rhizobia via increasing populations in large nodules (Figure 1). To examine differential rewards to more beneficial rhizobia in larger nodules, we focused on two *Medicago* hosts (A17 and R108) with intermediate selectivity but substantial differences in nodule size.

First, we characterized nodule morphologies and rhizobial populations to measure if there is a fitness benefit for a strain to be in a large nodule as compared to a small nodule (Figure 3a). Compared to small nodules, the large nodules from A17 were 7.4 times heavier (*P* < 0.01), had 1.2 times more lobes (*P* < 0.01), were 2.8 times longer (*P* < 0.001), and released approximately 6.4 times more rhizobia (*P* < 0.05). By contrast, for R108, large nodules were 13 times heavier, had 8.2 more lobes, were 2.5 times longer, and released approximately 14.4 times more viable rhizobia (all *P* < 0.001). The differences between A17 and R108 were particularly evident for large nodules; large nodules from R108 were heavier, had more lobes, and released far more rhizobia than large nodules from A17 (Figure 3b-e, Table S4). The larger differences in rhizobia released from large versus small in R108 than A17 suggest that while both hosts could enrich beneficial strains, R108 has more *potential* than A17 based on the larger asymmetry in nodule rhizobial population sizes.

**Figure 3.**
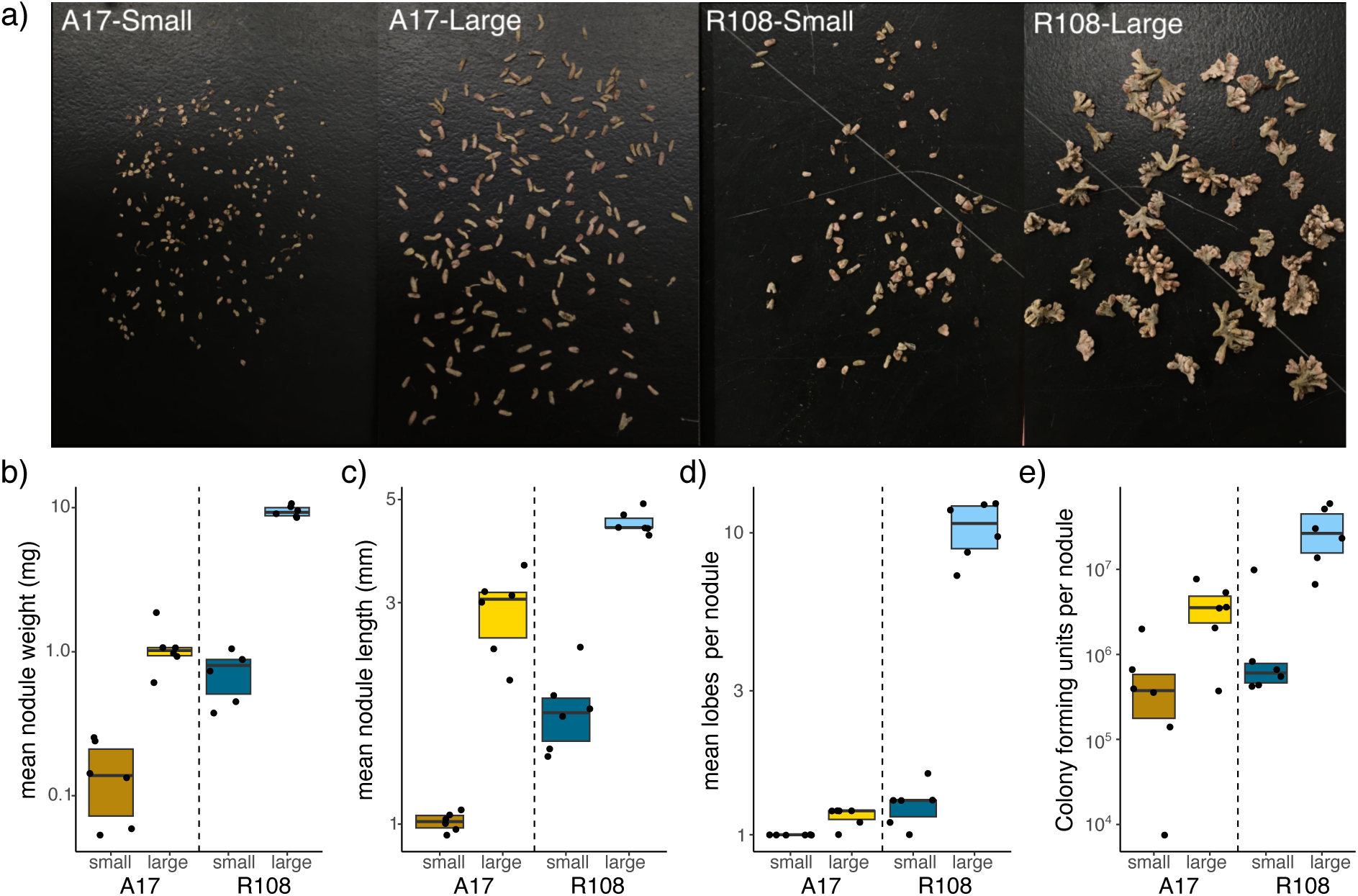
Nodule characteristics in two hosts, A17 and R108. a) Examples of single replicates of large and small nodule pools from our two host genotypes. (b-e) Nodule characteristics (weight, length, lobes, colony-forming units) of small and large nodules from A17 and R108 hosts. Each data point represents a replicate. The average weight per nodule is based on the fresh weight of the entire pool of nodules, and colony data is the average of replicate dilutions from haphazardly selected nodules from each class. All traits are on a log10 scale. All traits differed (p<0.05) between hosts and nodule size classes based on linear regressions and ANOVA (Table S4).

Second, for enrichment to occur, specific strains must be consistently enriched in large nodules compared to small nodules. We estimated the relative fitness of strains in the small and large nodule pools to measure the strength and consistency of host selection and the potential for host rewards. We expect 1) strains in small nodule pools to represent young and/or unrewarded nodules and 2) strains that derive more rewards from hosts to be overrepresented in larger nodules. If all nodules increase in size at the same rate regardless of strain identity, we would expect strain representation in small and large pools to be the same. A redundancy analysis (RDA) of strain relative fitness in large and small nodules in A17 and R108 revealed that host, size, and their interaction explained ∼42% of the variation in the composition of strain communities. Moreover, the interaction between Host and Size (*P* <0.001, df=1) suggests that hosts enriched different strains in large nodules. Subsetting the data by host reveals that nodule size explained 41% (*P* =0.003, df=1, Figure S7a) of the variation in strain communities in A17 and 17% in R108 (*P* =0.012, df=1, Figure S7b). Host identity was a strong determinant of representation in the nodule communities, and small and large nodules consistently supported different strain communities across replicates (Figure 4a, Table S5). Thus, we established the *potential* for adaptive host enrichment in both hosts.

**Figure 4:**
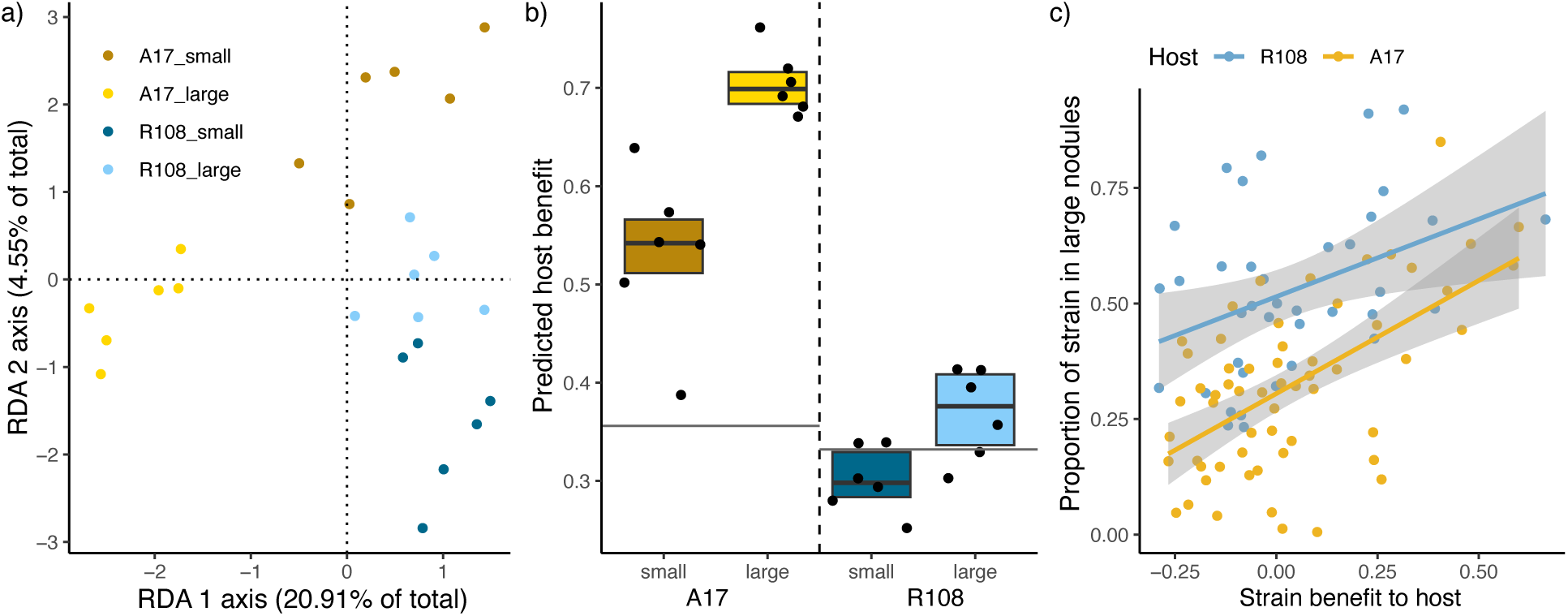
Adaptive strain enrichment in two model host genotypes. a) Multivariate redundancy analysis (RDA) of strain fitness in large and small nodule pools in A17 and R108 hosts revealed that host, nodule pool size, and their interaction account for approximately 41% of the variation in strain community composition. The interaction between host and size (*p*<.001, df=1) indicated that hosts enriched different strains in larger nodules. Note that constrained axis RDA 1, which differentiates large and small A17 nodules, explains 4-5 times more variation than constrained axis 2. b) The predicted benefit of the rhizobia found in nodules differs between hosts and nodule size (*p*=0.024, df=1). The predicted benefit is estimated by multiplying each strain’s benefit to hosts in a single-strain experiment by its frequency in the nodule pool and summing this across all strains. Since hosts vary in overall size, this sum is scaled such that one represents a nodule community composed entirely of the most beneficial strain, while zero signifies a community composed of the least beneficial strain. c) The relationship between the rewards rhizobia receive and the benefit those strains provide to plant hosts A17 and R108 during single-strain inoculations. The rewards received are measured by their proportional representation in large nodule pools versus the total representation in large and small pools. Each dot represents the median strain proportion across six replicate pots. Single-strain plant benefit predicted the enrichment of a strain in larger nodules for both hosts, but explained more variation in A17 (34%) than in R108 (15%). Benefits for b and c were estimated by measuring plant biomass in a single-strain inoculation experiment under nitrogen-free conditions (See Tables S4-S6 for complete statistical results).

Third, we examined strain communities in small nodules to assess the potential for pre-nodulation partner choice mechanisms to explain rhizobial fitness. Strains in small nodule pools are a combination of both young and unrewarded nodules. We found that small nodules support rhizobial communities that are predicted to be more beneficial than expected by chance associations for A17 but not R108; strains in small A17 nodules extract 55% of the potential benefit in the initial inoculum as compared to 35% expected by chance, while strains in small R108 nodules are not more beneficial to R108 than expected by chance 30% vs 33% (Figure 4b, Table S6). These results suggest pre-nodulation partner choice mechanisms can contribute to rhizobial strain success, at least for one host genotype.

Lastly, we focused on selection mechanisms occurring after nodules form by comparing the host benefits of strains in large and small pools. We took two complementary approaches to test whether hosts could adaptively enrich strains after nodules formed (Figure 1). Consistent with the preferential enrichment of beneficial strains, particularly by A17, the estimated benefit of the nodule community increased by ∼ 20% in large nodules than in small nodules in A17 but < 10% in R108 (Figure 4b, both *P* < 0.01, *P*_host*size_ = 0.018). We then examined the relationship between strain enrichment in large nodules and the benefit plants obtained from each strain. Consistent with hosts rewarding more beneficial strains, this relationship was significantly positive for both hosts (Figure 4c, Table S7, r = 0.61, *P* < 0.0001). However, it was weaker for the less selective host R108 (Figure 4c, r = 0.3, *P* = 0.07). Overall, our results suggest A17 was better than R108 at enriching the fitness of more beneficial strains.

## DISCUSSION

Mutualisms are expected to break down unless mutualistic partners limit the evolution of genotypes that reap benefits but do not reward mutualistic partners (Sachs and Simms, 2006; Heath and Tiffin, 2009; Jones et al., 2015). Here, we show ample genetic variation in host selectivity in a model legume species (Henry and Nevo, 2014). We identify candidate genes contributing to this variation to garner clues about potential mechanisms. The annotated functions of these genes point towards the importance of symbiont access to Zinc/Iron, cell wall modification, DNA methylation, and under-characterized plant hormones. Furthermore, we demonstrated an association between nodule morphology and selectivity: on average, host accessions with shorter nodules tend to be more selective. To ask whether hosts differ in their ability to enrich beneficial rhizobia adaptively is much more challenging, so we focused on two host accessions with contrasting nodule morphology, A17 and R108. While both *Medicago* accessions were able to preferentially enrich the fitness of more beneficial symbionts by increasing the size of the nodules they inhabited, contrary to our expectation, the A17 host with shorter nodules and smaller variance in nodule size was more effective at preferentially rewarding beneficial strains via both pre- and post-nodule formation mechanisms. Our results suggest the importance and feasibility of leveraging natural genetic variation in hosts and symbionts to decipher the mechanisms underlying mutualism maintenance (Henry and Bergelson, 2023).

Host selectivity is sometimes inferred directly from bacterial diversity in nodules (Epstein et al., 2023). This whole-plant diversity may be appropriate when querying ecological and evolutionary shifts in rhizobia communities. However, when identifying mechanisms that affect rhizobial diversity in nodules, host sampling processes can be confounding (e.g., hosts that form more nodules may exhibit increased diversity due to increased sampling, rather than host control). After accounting for among-host variation in nodule number, we uncovered heritable variation in host selectivity within *Medicago.* While most host genotypes were weakly selective, a handful were highly selective, suggesting the potential for selecting on this trait (Clouse and Wagner, 2021; Henry and Bergelson, 2023; Porter et al., 2024).

Moreover, we identified allelic variation associated with host selectivity. These candidates differ from those highlighted in the diversity-based GWAS in Epstein et al. 2023, suggesting that we are focusing on a different trait. The top candidates point to interesting host genes, including MtLegin27, a leginsulin-related MtN11/16/17 family gene(de Bang et al., 2017; Boschiero et al., 2020). Not much is known about these secreted hormone-like cysteine-rich peptides, but three similar genes (Legin47, 31, 42) are upregulated 72 hours post-inoculation with rhizobia (Gao et al., 2022). Another promising candidate, MtZIP11 - Zinc-Iron Permease 11 - is highly expressed in nodules and is a close homolog of MtZIP6, which specifically transports Zinc into rhizobia-infected host cells (Abreu et al., 2017). The association with host selectivity is particularly interesting as Zinc has recently been shown to enable the function of FUN (FIXATION UNDER NITRATE), a transcription factor regulating nodule senescence and nitrogen fixation in response to nitrate availability in *Lotus japonicus* (Lin et al., 2024). The GWAS also identified a gene (MtrunA17Chr4g0058051) encoding a putative pectate lyase (Xie et al., 2012; Liu et al., 2019; Su, 2023; Zhang et al., 2024) that increases in expression within the first three days after exposure to rhizobia (Pereira et al., 2024). MtMATE38 is a member of a mostly uncharacterized family of 70 multidrug & toxic compound extrusion transporters that are likely biologically important (e.g., transporting citric acid, flavonoids, and aluminum) (Wang et al., 2017). Finally, we identified another gene whose expression differs in early nodule development (MtrunA17Chr3g0127911), annotated as a putative transcription initiation Spt4, spt4/RpoE2 zinc finger (Pereira et al., 2024). Similar transcription factors are involved in RNA-directed methylation in *Arabidopsis* (Köllen et al., 2015). Together, these candidates point to potential mechanisms of host selectivity in *Medicago* beyond the frequently studied NCR peptides (de Bang et al., 2017; Montiel et al., 2017; Horváth et al., 2023). Figuring out how plants enrich beneficial bacteria is an area of active exploration (Henry and Bergelson, 2023) and may lead to exciting mechanisms for stabilizing mutualisms in agricultural systems (Denison et al., 2003; Kiers and Denison, 2014; Busby et al., 2017).

Associating nodule morphological characteristics with host selectivity could point us towards developmental mechanisms underlying that selectivity. Multiple significant relationships between suites of correlated nodule morphology traits and host selectivity suggest that hosts that form numerous nodules that are shorter, smaller, less elongated, and with proportionally fewer lobed nodules tend to be more selective on average. However, we did find that the single best predictor of host selectivity was the average major axis length (i.e., shorter nodule length on average), suggesting this trait could be used as a proxy in future studies for the primary suite of nodule morphology differences. While this trait explained a relatively modest amount of variation (12%), our results suggest that mechanisms by which hosts can control the size of symbiotic organs may be important, especially as recent research suggests that rhizobial bacteria can control nodule size via expression of plant hormones(Nett et al., 2022). More broadly, this research serves as a starting point for connecting organ form to function in symbiosis research, suggesting that expanding research in this direction would be worthwhile.

Increased host selectivity can be adaptive or not, and figuring out how plants can or cannot preferentially enrich beneficial bacteria while repelling pathogens or cheaters is an area of active exploration (Henry and Bergelson, 2023). Although both host genotypes we tested were able to enrich beneficial rhizobia strains, our results make clear that the preferential rewarding of more beneficial symbionts is far from perfect, and both pre- and post-nodulation processes affect strain frequencies in nodules. This variation in strain-specific rewards is important as it may allow less beneficial strains to reap the reproductive benefits of symbiosis, thereby contributing to maintaining mediocre rhizobia strains in populations (Bever et al., 2009; Bever, 2015). More generally, our results suggest pairing assessments of single-strain benefits with assessments of nodule strain competitive fitness can identify host genotypes that best maintain mutualistic rhizobia populations in the face of diverse rhizobial populations in natural and agricultural soils (Mendoza-Suárez et al., 2020; Batstone et al., 2022; Burghardt and diCenzo, 2023).

## ACKNOWLEDGEMENTS

This work was supported by the National Science Foundation grants IOS-1724993, 1856744, 2243817, and 2243819 to Burghardt and Tiffin and completed through computing resources provided by the Minnesota Supercomputing Institute (MSI) at the University of Minnesota and sequencing support from the Department of Energy Community Sequencing Project #503446. LTB was partly supported by USDA NIFA Grant 2022-67013-36860 and this work is/was supported by the USDA National Institute of Food and Agriculture and Hatch Appropriations under Project #PEN04760 and Accession #1025611. We thank Adam Kostanecki for taking the nodule pictures, members of the Tiffin and Heath Labs for helping with the harvesting, and Burghardt Lab members for feedback that improved the manuscript.

## DATA AVAILABILITY

All code, data sets, and results are available on the Burghardt Lab Github Repository

## AUTHOR CONTRIBUTIONS

LTB and PT conceived the study, analyzed data, created figures, and wrote and revised the paper. LTB implemented the experiment. BE/PS/JS performed bioinformatic and statistical analyses, drafted text and figures, and edited the manuscript.

---

**Table S1:**
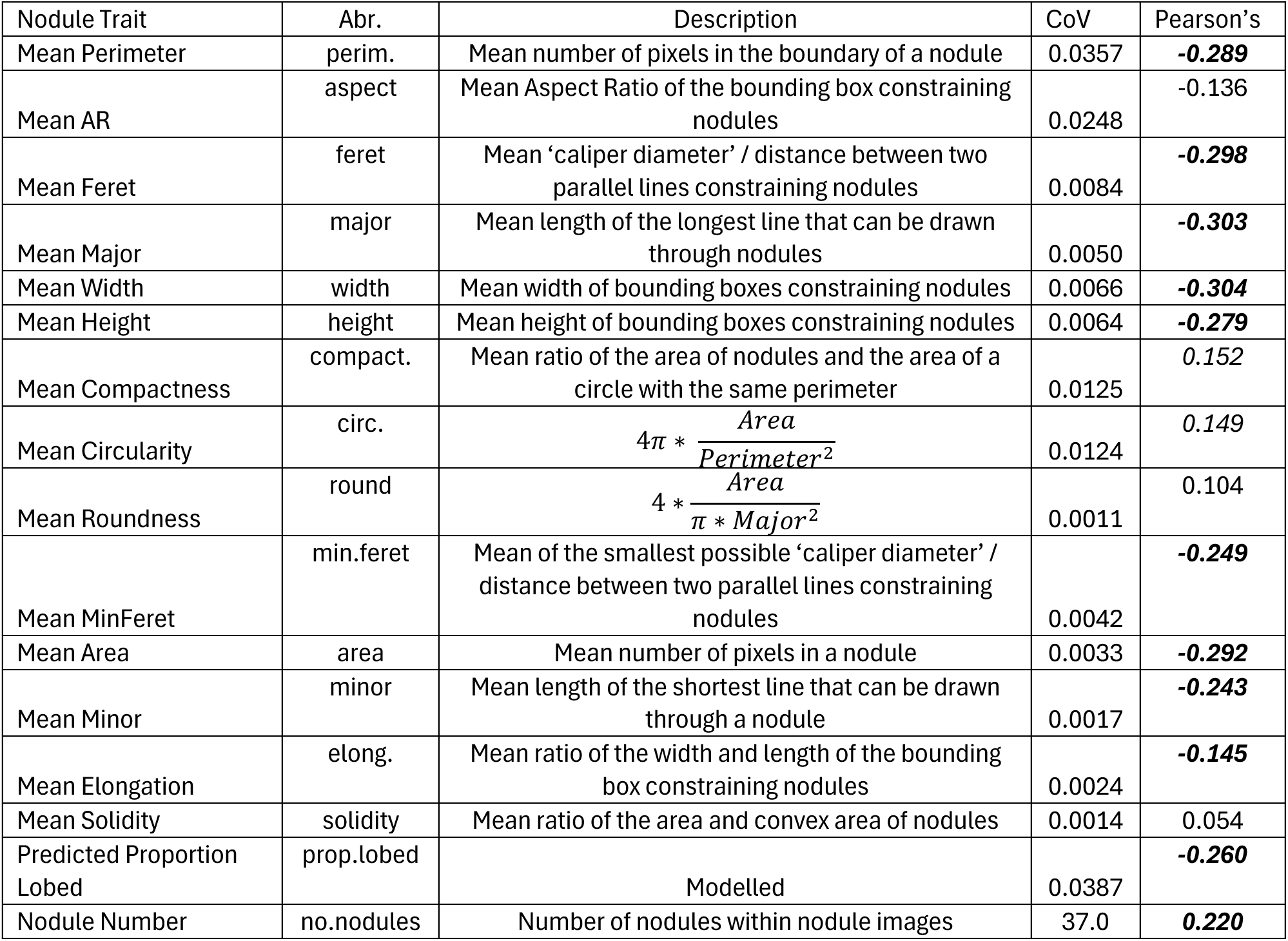
Nodule traits measured from 4-5 replicate pots of 202 Medicago accessions inoculated with 89 strains of *Sinorhizobium meliloti* (Epstein et al. 2023). Coefficient of Variation (CoV) for each trait and Pearson’s r describing the trait correlation with Host Selectivity (*p < 0.05 in italics*) and Traits that are significant at ***p < 0.05*** after correction for multiple test are ***in bold italics***).

**Table S2:** Top 10 candidate SNP variants for host selectivity selected based on *p*-values from likelihood ratio tests. MAF is the Minor allele frequency. Beta is the effect size from the genome-wide association and reflects that non-A17 alleles tend to increase host selectivity.

| ChromosomeID-Position-BaseChange | Chr | -<br>log<sub>10</sub>($p$) | MAF | beta | Closest gene in the Medicago V5 genome |
| --- | --- | --- | --- | --- | --- |
| CM010654.1-26952852-C | 7 | 9.365 | 0.05 | 0.148 | MtrunA17_Chr7g0236791- MtLegin27- Leginsulin |
| CM010650.1-51530113-A | 3 | 8.299 | 0.06 | 0.124 | MtrunA17_Chr3g0135691- MtZIP11- Zinc-Iron Permease 11 |
| CM010651.1-51084261-T | 4 | 7.973 | 0.31 | 0.111 | MtrunA17_Chr4g0058051- putative pectate lyase |
| CM010653.1-25128406-A | 6 | 7.660 | 0.05 | 0.156 | MtrunA17_Chr6g0472811- snoR71-ncRNA |
| CM010651.1-51080330-T | 4 | 7.586 | 0.26 | 0.112 | MtrunA17_Chr4g0058051- putative pectate lyase |
| CM010654.1-23572058-T | 7 | 7.563 | 0.06 | 0.114 | MtrunA17_Chr7g0233111- putative protein |
| CM010653.1-32706149-T | 6 | 7.556 | 0.08 | 0.105 | MtrunA17_Chr6g0477961- MtMATE38 -multidrug & toxic compound extrusion family |
| CM010650.1-45833295-C | 3 | 7.405 | 0.06 | 0.129 | MtrunA17_Chr3g0127911- Putative transcription initiation Spt4, spt4/RpoE2 zinc finger |
| CM010649.1-16046163-T | 2 | 7.329 | 0.12 | 0.093 | MtrunA17_Chr2g0298091- putative transferase |
| CM010654.1-23572805-T | 7 | 7.046 | 0.07 | 0.108 | MtrunA17_Chr7g0233111- putative protein |

**Table S3:**
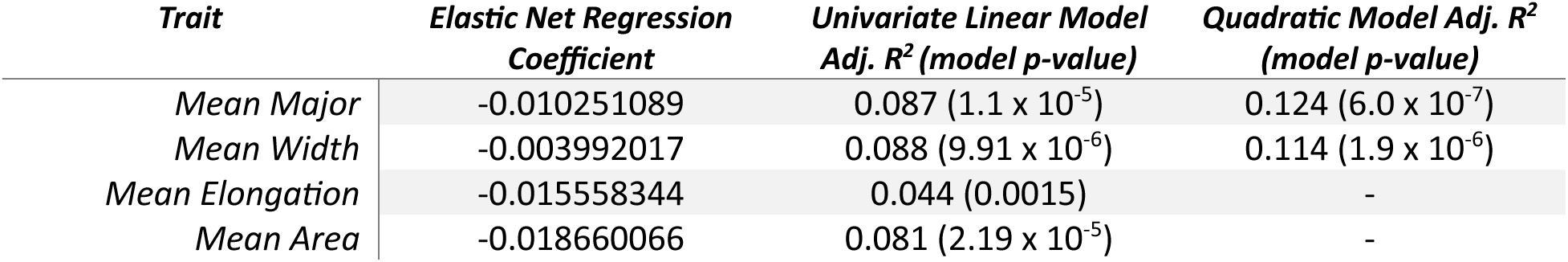
Host Selectivity x Nodule Trait model summaries of four traits selected via Elastic Net Regression including all nodule traits as potential predictors.

| Trait | Elastic Net Regression<br>Coefficient | Univariate Linear Model<br>Adj. R<sup>2</sup> (model $p$-value) | Quadratic Model Adj. R<sup>2</sup><br>(model $p$-value) |
| --- | --- | --- | --- |
| Mean Major | -0.010251089 | 0.087 (1.1 x 10<sup>-5</sup>) | 0.124 (6.0 x 10<sup>-7</sup>) |
| Mean Width | -0.003992017 | 0.088 (9.91 x 10<sup>-6</sup>) | 0.114 (1.9 x 10<sup>-6</sup>) |
| Mean Elongation | -0.015558344 | 0.044 (0.0015) | - |
| Mean Area | -0.018660066 | 0.081 (2.19 x 10<sup>-5</sup>) | - |

**Table S4:**
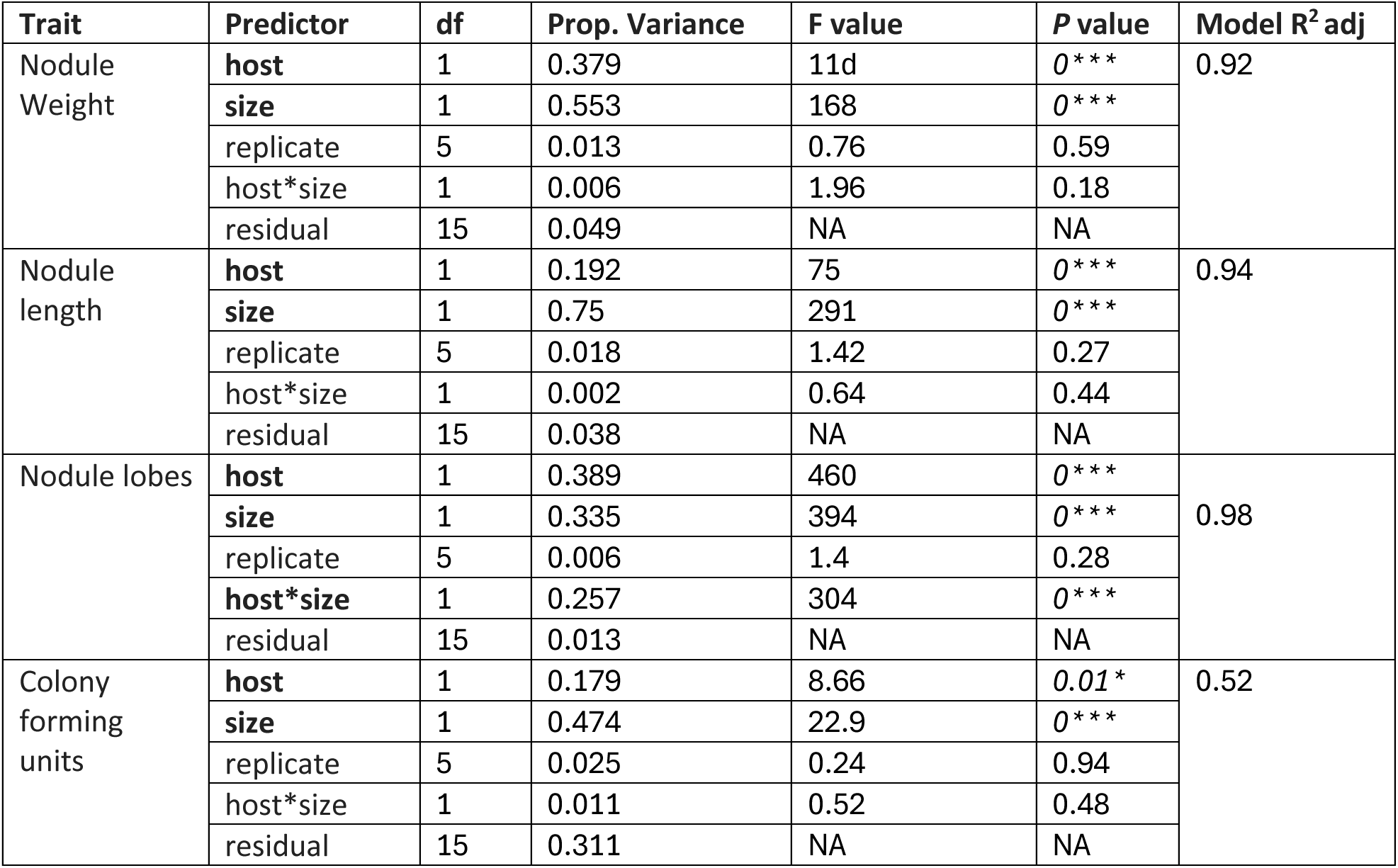
ANOVA results for linear models of individual nodule traits from sequenced pools (Figure 3).

**Table S5:** PERMANOVA results from RDA analysis of strain communities in nodules pools (Figures 4a and S5).

| Models | predictor | df | Prop Variance | F value | P value | R<sup>2</sup> adj |
| --- | --- | --- | --- | --- | --- | --- |
| Both Hosts | host | 1 | 0.18 | 6.10 | 0.001** | 0.34 |
|  | size | 1 | 0.13 | 4.56 | 0.001** |  |
|  | Host*size | 1 | 0.12 | 4.03 | 0.002** |  |
|  | residual | 20 | 0.58 | NA | NA |  |
| A17 only | size | 1 | 0.41 | 6.98 | 0.004** | 0.35 |
|  | residual | 10 | 0.59 | NA | NA |  |
| R108 only | size | 1 | 0.17 | 2.02 | 0.016* | 0.085 |
|  | residual | 10 | 0.83 | NA | NA |  |

**Table S6:**
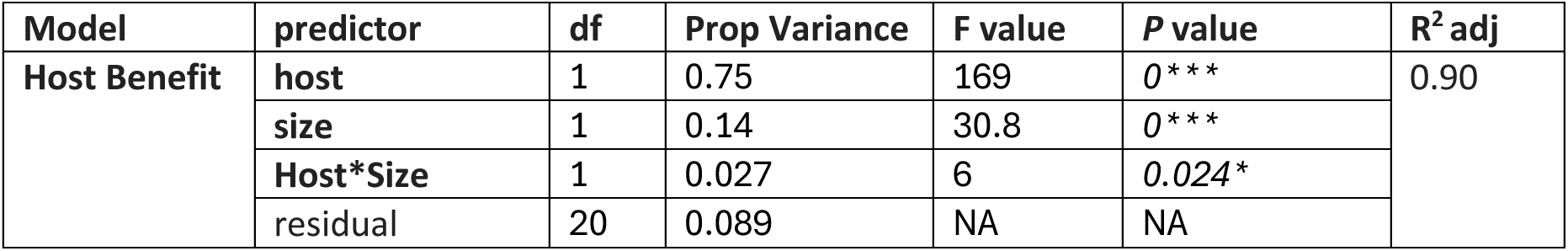
Statistical results for predicted host benefit of each strain community inferred from previous single-strain experiments (. **Figure 4b)**

| Model | predictor | df | Prop Variance | F value | P value | R<sup>2</sup> adj |
| --- | --- | --- | --- | --- | --- | --- |
| Host Benefit | host | 1 | 0.75 | 169 | 0*** | 0.90 |
|  | size | 1 | 0.14 | 30.8 | 0*** |  |
|  | Host*Size | 1 | 0.027 | 6 | 0.024* |  |
|  | residual | 20 | 0.089 | NA | NA |  |

**Table S7:** ANOVA results from linear models of strain relative representation in large vs. small nodules (. **Figure 4c).**

| Model | predictor | df | Prop Variance | F value | P value | R<sup>2</sup> adj |
| --- | --- | --- | --- | --- | --- | --- |
| Both Hosts | host | 1 | 0.20 | 33.5 | 0*** | 0.42 |
|  | benefit | 1 | 0.23 | 38.2 | 0*** |  |
|  | Host*benefit | 1 | 0.006 | 0.097 | 0.32 |  |
|  | residual | 93 | 0.56 | NA | NA |  |
| A17 only | benefit | 1 | 0.34 | 29.7 | 0*** | 0.33 |
|  | residual | 57 | 0.66 | NA | NA |  |
| R108 only | benefit | 1 | 0.15 | 6.4 | .016* | 0.13 |
|  | residual | 36 | 0.85 | NA | NA |  |

**Figure S1:**
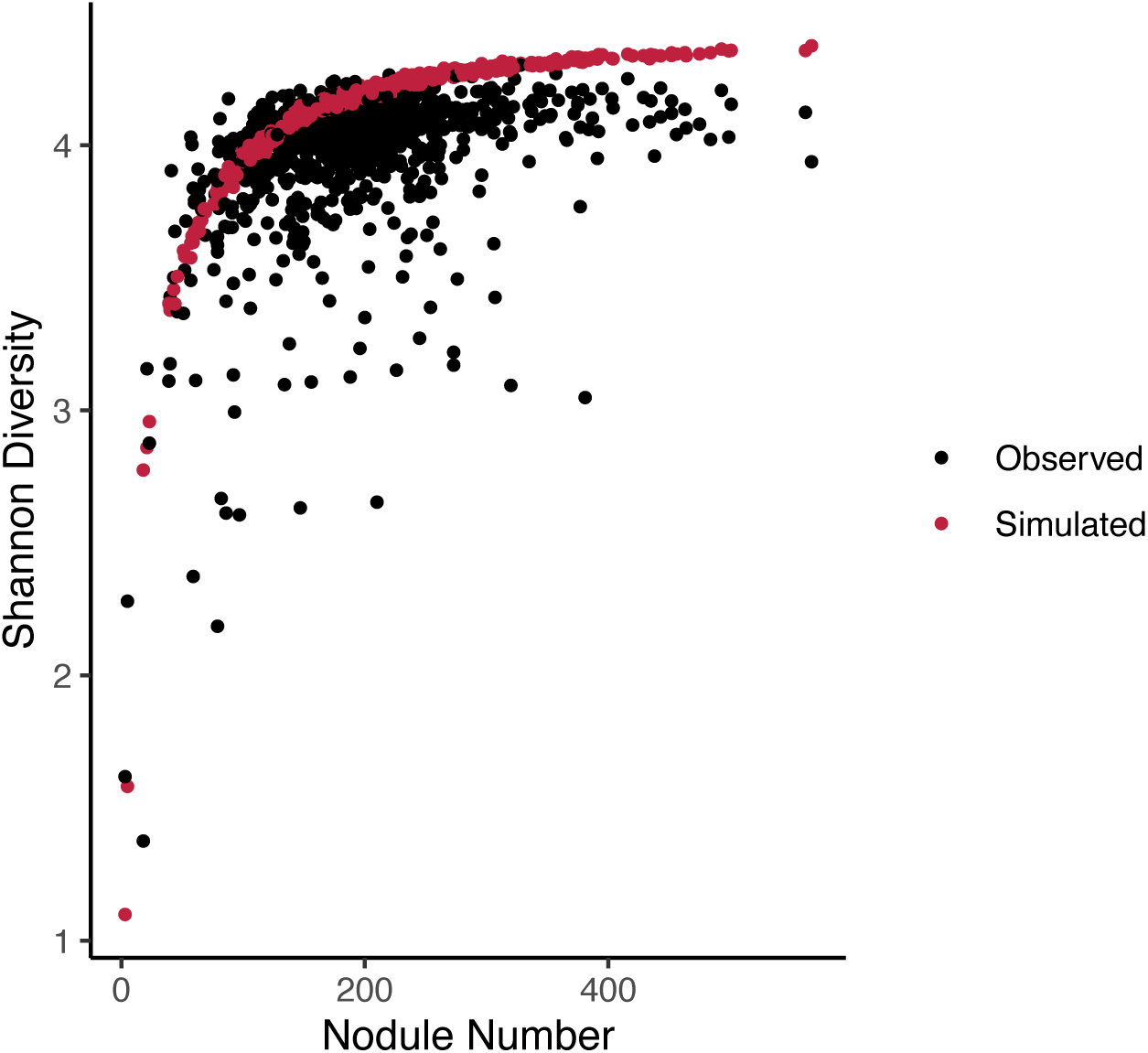
Relationship between nodule number and Shannon diversity for simulations and empirical data. Red dots are the average of 10 simulations of neutral nodule assembly at each empirically observed nodule number (red dot). Black dots represent empirical data on nodule Shannon diversity estimated for each of the four replicate pots of 202 Medicago accessions. ‘Host selectivity’ is estimated by subtracting the observed value from the simulated value for a given nodule number.

**Figure S2:**
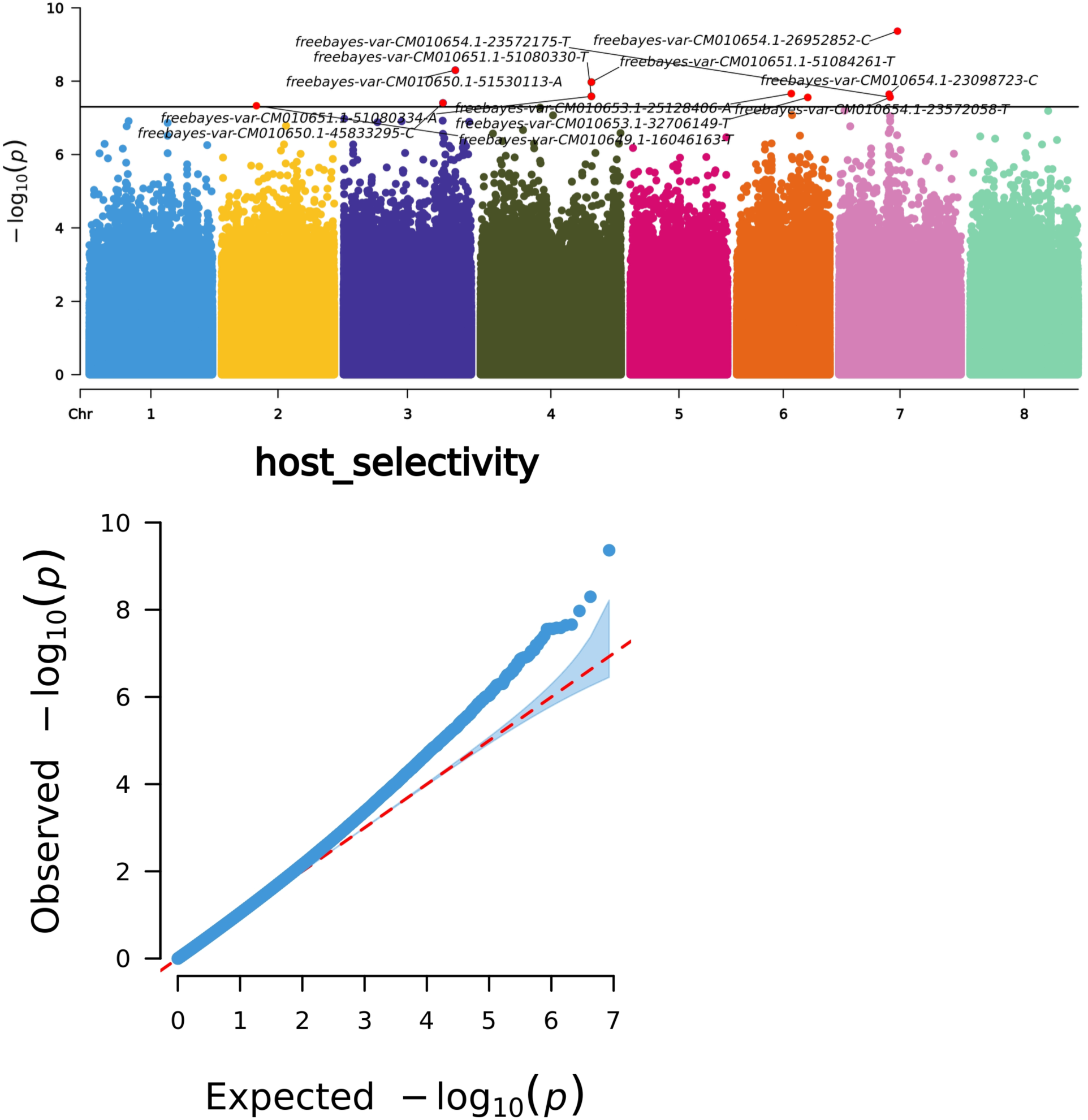
Visualization of results from the genome-wide association analysis of the host selectivity traits Manhattan plot (top) showing the 10 candidate variants with *p* values less than 1×10^-7^. QQ plot (bottom) of expected vs. observed p-value distribution. Candidate information is in Table S2.

**Figure S3:**
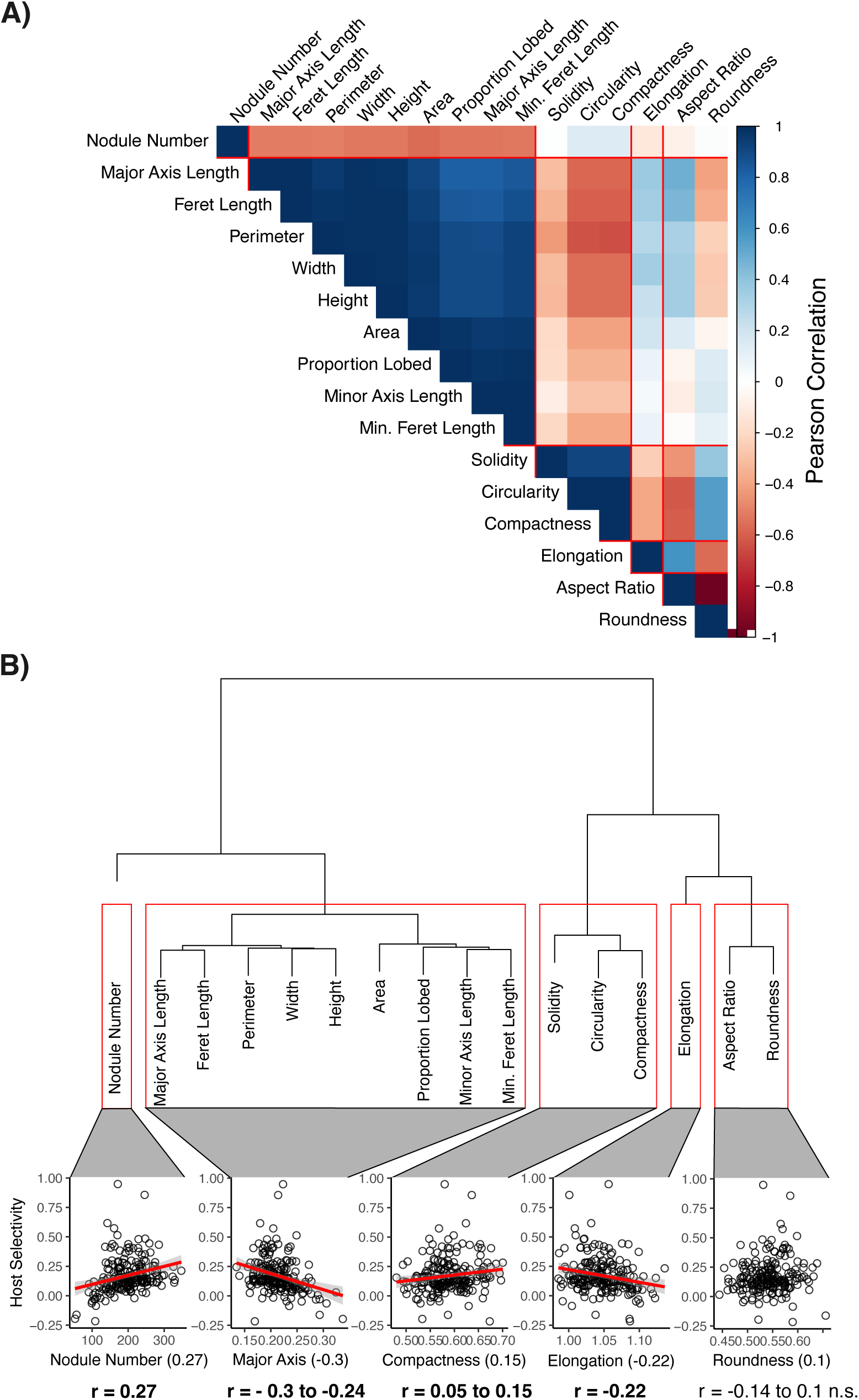
Nodule morphological traits cluster into five major groups at > 0.80 (a). Visual summary of linear regression model with one representative trait from each of the five clusters with the host selectivity trait (b). Significant relationships include a red line.

**Figure S4:**
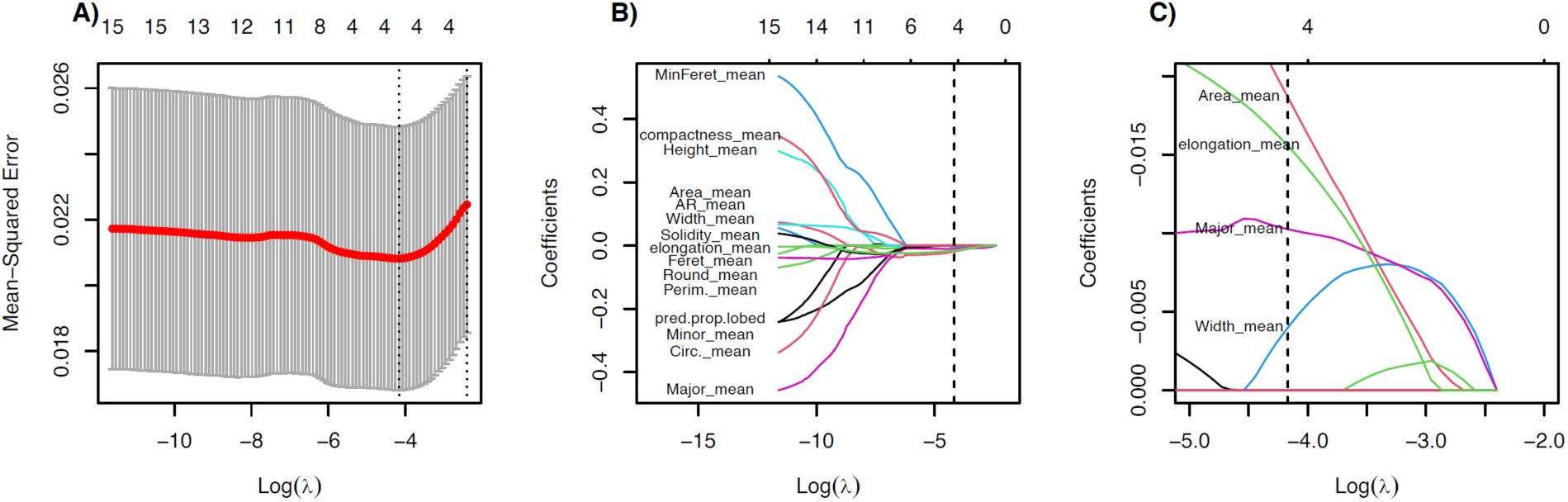
Summary of Host Selectivity elastic net regression. The size of the λ penalty parameter selected by cross-validation (n folds = 10) through comparison of Mean-Squared Error (λ = 0.155; Log(λ) = −4.17). Declining coefficients of nodule trait parameters across increases in the shrinkage penalty parameter λ (B). Focused comparison of non-zero nodule trait coefficients selected by elastic net regression at Log(λ) = −4.17 (C).

**Figure S5:**
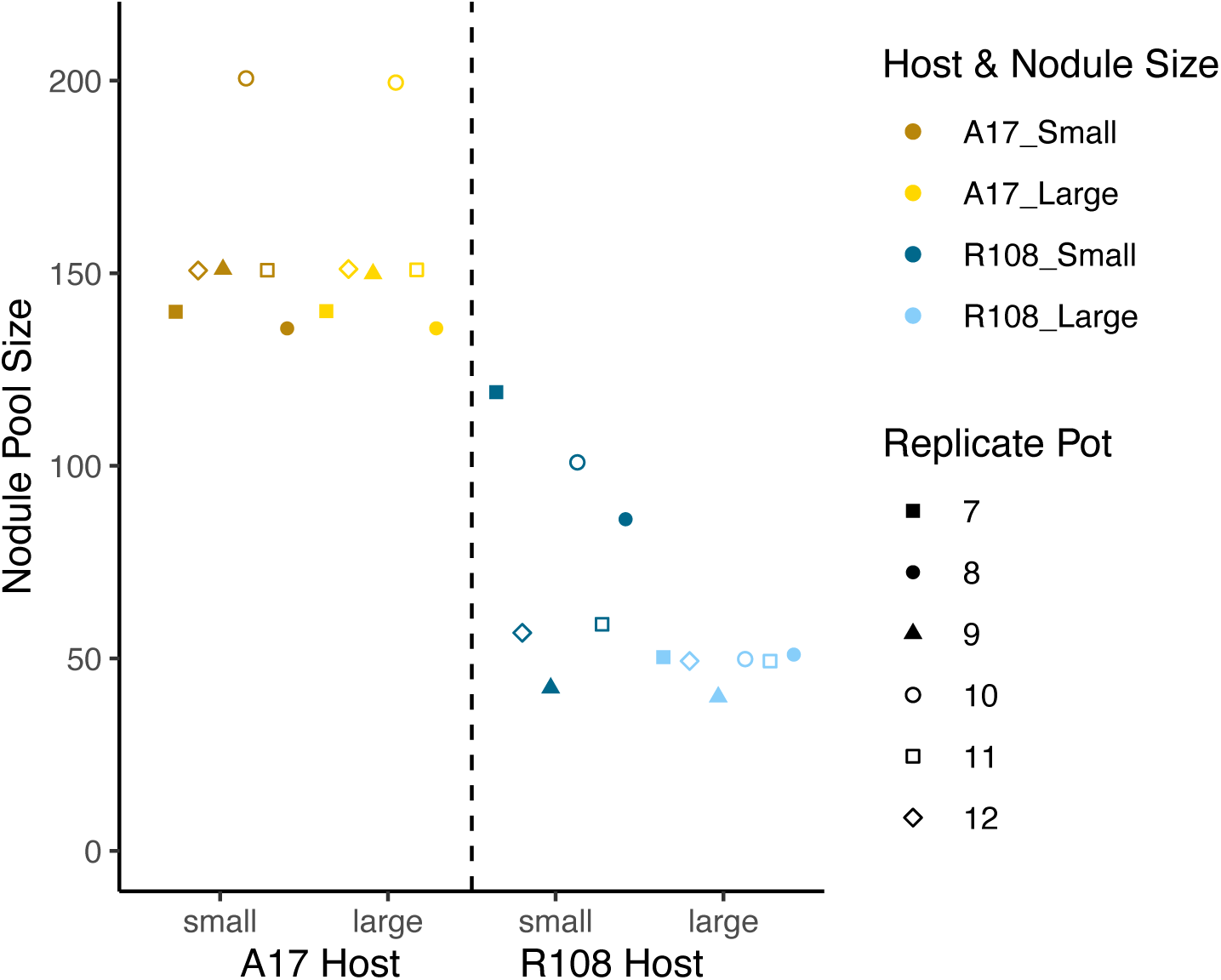
Size of Sequenced Nodule Pools. A17 nodules have identical nodule pool sizes, whereas 3 of the R108 pools had smaller nodules in the large pool due to processing volume and nodule size limitations. However, this did not affect Shannon diversity (Figure S6), suggesting nodule pools were saturated.

**Figure S6:**
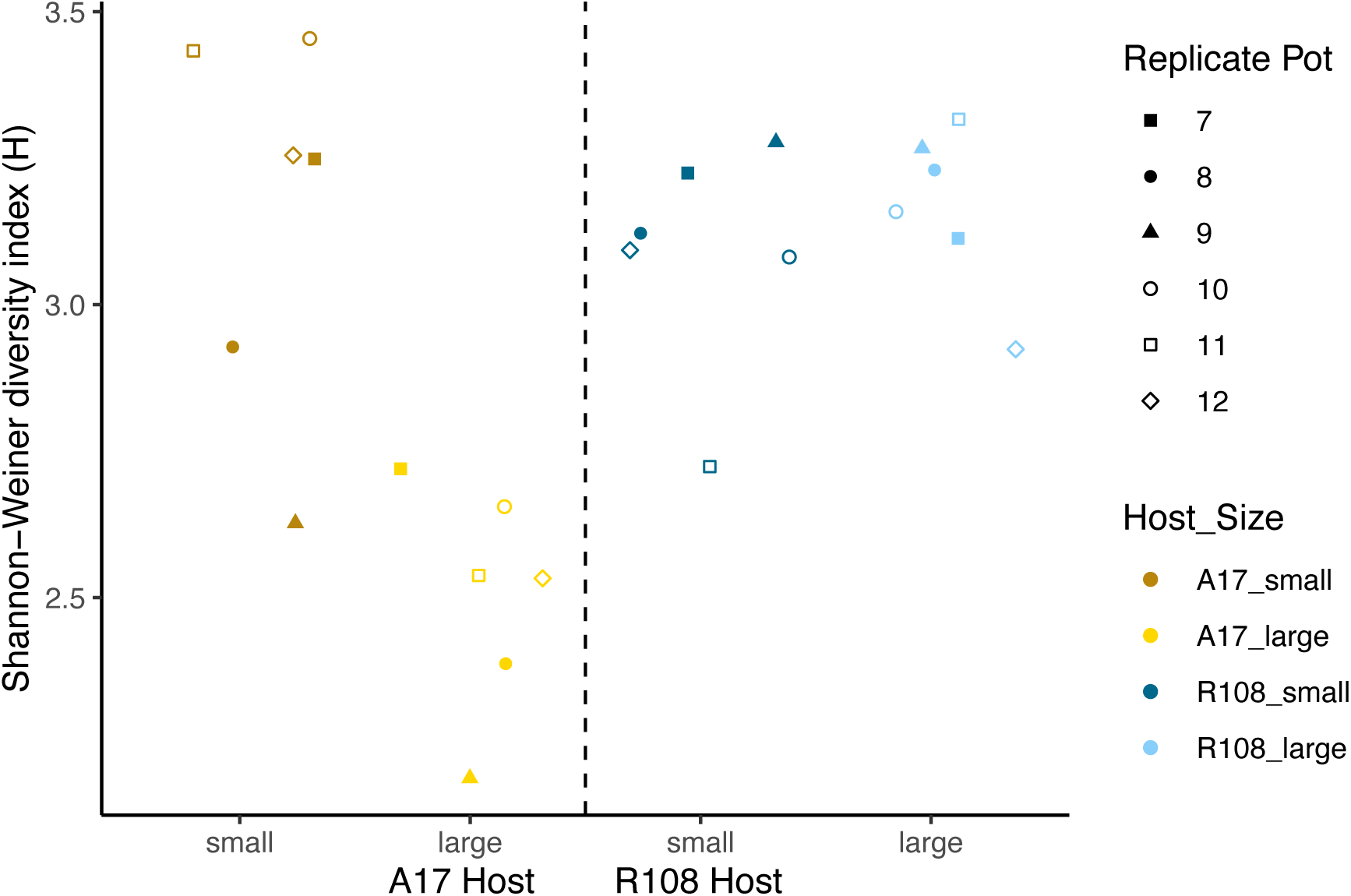
Strain diversity in Sequenced Nodule Pools. Note that differences in nodule pool size in Figure S5 for pots 7, 8, and 10 do not alter strain diversity patterns. This suggests that the practical size limitation in half of the large nodule pools for R108 is not affecting our inferences.

**Figure S7:**
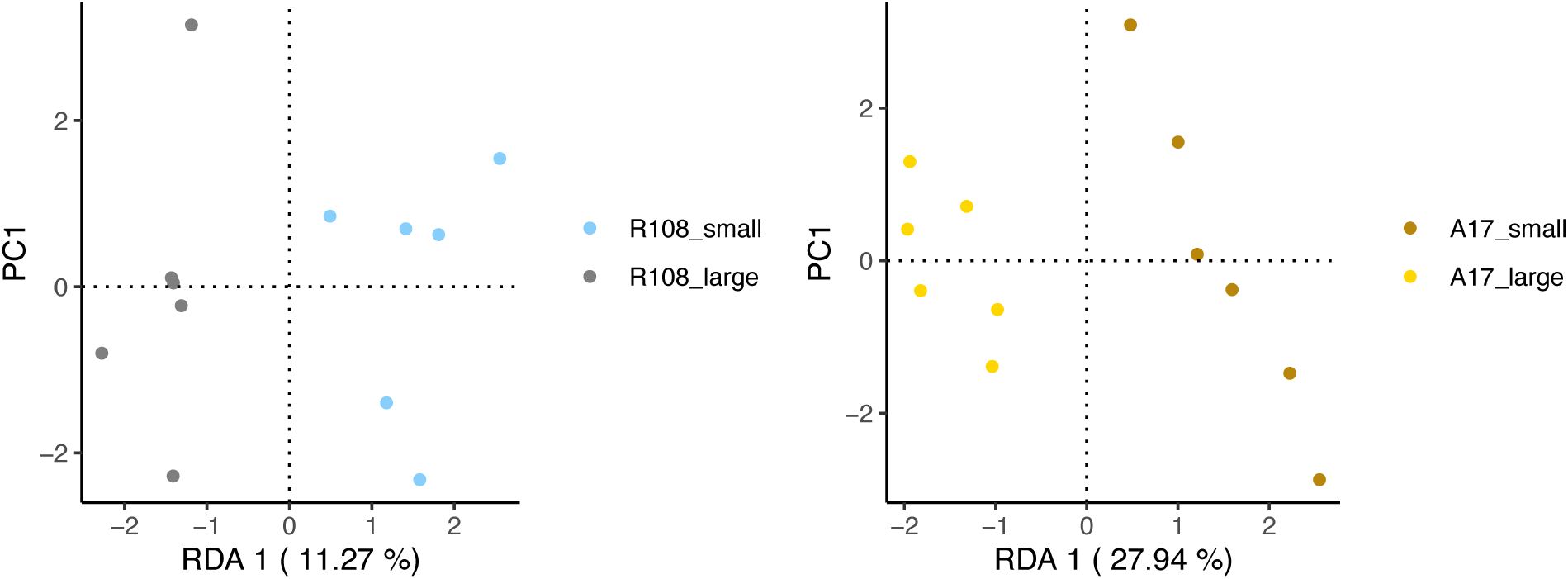
RDA analysis conducted independently for large and small nodule strain communities in A17 (left) and R108 (right). See Table S7 for statistical results.

